# Can a Tiny Brain Plan? Reconstructing a Predatory Detour in the Jumping Spider *Portia labiata*

**DOI:** 10.64898/2026.11.04.763773

**Authors:** Finn J. Petrie, Min Tan, Chen Li, Ximena J. Nelson, Daiqin Li, Malcolm A. MacIver

## Abstract

Planning is the process of evaluating alternative futures to predict outcomes and guide action. Evidence for this process exists in several vertebrates, but remains contested in invertebrates. Jumping spiders in the genus *Portia* show planning-like behavior in lab studies, but how this translates to the wild is unclear, where hunts of potentially lethal spider prey occur over durations of more than an hour in dense foliage. We present the first fully video-recorded wild hunt of any *Portia* species, with a 3D reconstruction of its forty-two minute long route to attack an orb-weaving spider. The reconstruction suggests *P. labiata* may have planned its approach, avoiding a route roughly half as long but potentially more conspicuous to the prey’s sensory systems.

## Introduction

Planning, the evaluation of alternative futures to guide action, is thought to require a large brain and complex neural structures, and evidence for it comes almost entirely from mammals and birds^1–3^. *Portia*, a genus of jumping spiders (Salticidae) native to Asia, Australasia, and Africa^4^, shows evidence of planning-like behavior despite having a brain orders of magnitude smaller than those of vertebrates^5–7^, raising the possibility that planning may not be limited to a larger brain or complex structures such as prefrontal cortex^8^. Arthropods and verte-brates diverged more than 600 million years ago^9, 10^ from a small marine worm without image-forming eyes^11, 12^, an ancestor unlikely to have planned. If *Portia* does plan, this would be a case of convergent evolution that could help illuminate what is necessary versus contingent in planning circuits across all animals.

Convergence would also open a question that mechanistic accounts of planning have largely set aside^1, 13^: what is the function of planning? The conditions under which planning pays off constrain what its neural machinery must compute, and if planning evolved independently in spiders and verte-brates, the conditions the two lineages share will highlight the types of situations under which planning pays off.

One hypothesis is that planning was selected for in the context of long-range visual detection of high-stakes goals in spatially complex environments (the Buena Vista hypothesis^14–19^). *Portia*’s hunting grounds appear spatially complex, so whether it may fit this hypothesis partially turns on how far it can see, which depends largely on visual acuity: how many cycles of black-to-white variation can be resolved in a single degree of the visual field (cycles per degree, cpd). Higher cpd extends the distance at which an object can be resolved. *Portia fimbriata*’s visual acuity has been measured anatomically at 12.5 cpd^20^ in a body with a mass of *≈* 50 mg. That is similar to a pigeon’s acuity in a body with a mass of *≈* 300 g^21, 22^, a 6,000-fold difference in body mass. For an arthropod comparison, *P. fimbriata*’s acuity is six times the highest acuity measured in any insect or non-salticid arachnid, the dragonfly *Anax junius* (*≈* 2 cpd)^21, 23, 24^; other spiders are far lower (e.g., *≈* 0.5 cpd in *Cupiennius salei* ^25^).

With its long-range vision in complex spatial domains, *Portia* has the ability to see its prey from far away, where many possible paths exist that are likely to vary in their probability of success—conditions that favor planning. Indeed, the presence of planning in *Portia* has been suggested, in part due to its detouring behavior, a complex strategy that involves losing sight of the prey and calculating a more effective route. Detouring, and arguably planning as a higher cognitive function, requires that an organism represent parts of its environment beyond its immediate surroundings, such as the location of a goal that is no longer in view^5, 7, 26^, and therefore may need a cognitive map. *Portia*’s detours in the wild can be particularly elaborate, with spiders losing sight of their prey for a median of 17 min, and a maximum of 81 min^27^.

Another signal that *Portia* may be capable of planning is the flexibility of its aggressive mimicry, a form of hunting where a predator resembles something harmless or beneficial to help it approach or attract prey^28^. *Portia* can manipulate its prey’s web in order to ‘control’ the prey’s behavior. For example, *Portia* will vibrate a web, perhaps in the same pattern as an ensnared insect, with the frequent result of luring the prey spider out, whereupon *Portia* will attack it^29–32^.

However, *Portia*’s ability to plan has primarily been studied in the lab, using artificial or dead lures. It is unclear whether planning pays off in the wild, where *Portia* must choose among many routes through clutter toward prey that can detect it and fight back. Among jumping spiders, *Portia* is unusual in specializing in preying on other spiders that can injure or kill it^33–35^. Perhaps as a result, hunts are often abandoned, sometimes hours after their initiation^27, 33^, making field data unusually hard to collect, in addition to the challenge of recording in dense vegetation. Two professional natural-history film crews, each spending more than a week in the field with one of us (D.L.), did not succeed in recording a complete wild hunt.

Related to these issues, while there have been detailed descriptions of wild hunts in *Portia*^27^, this is the first wild hunt of any *Portia* species fully captured on video. After a failed attempt to lure the prey from its web, *P. labiata* appears to plan, evaluating alternative routes and rejecting a shorter silk-obstructed path in favor of a longer, unobstructed route enabling a novel attack from beneath the web.

## Results

The hunt was recorded on August 1, 2024 in the Cameron Highlands in Malaysia. A meter away from the hunt location, a female *P. labiata* was found, and was moved to near where we observed a juvenile orb-weaver prey spider (*Argiope versicolor*) on its web. The data from the hunt included videos from two handheld smartphones and a high resolution laser scan of the environment (details: Methods).

The hunt lasted 41:34 (MM:SS) (Video SM1)^1^. As outlined in Fig. 1, *P. labiata* expressed multiple behaviors during the hunt—aggressive mimicry, rerouting after failure of that strategy, selection of a path that may have been vibrationally and visually inconspicuous to its prey, a stalking sequence, and the final attack from under the prey web’s discoid decoration which may visually shield *Portia* from the prey.

**Fig. 1:**
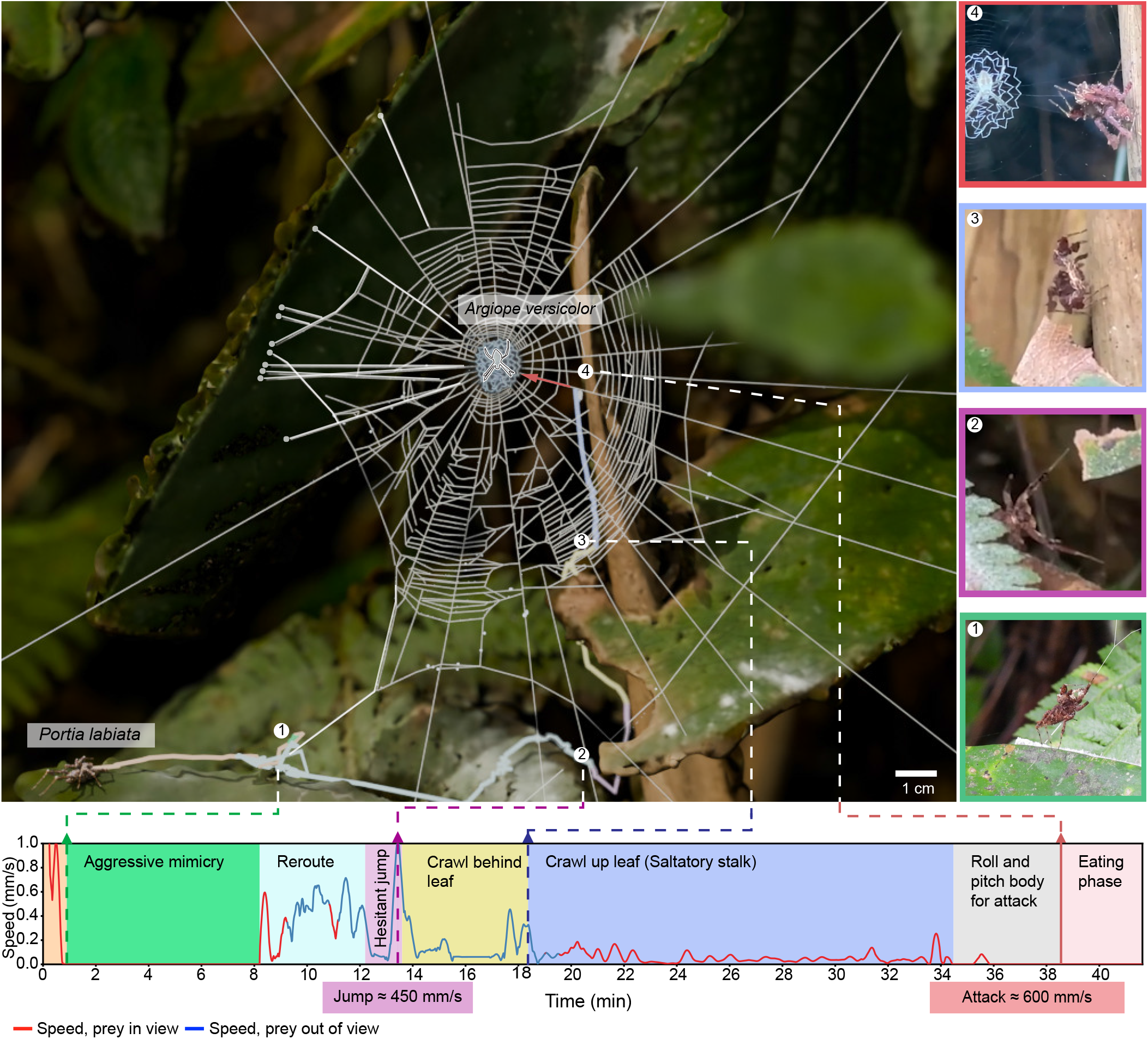
Timeline and trajectory of the hunt. The view shown here is from an interactive browser-based 3D reconstruction that allows moving through the space and time of the hunt (Hosted by GitHub: SA1). Guylines that connect to the left leaf are rendered thicker to highlight them, as well as one guyline used by *P. labiata* during putative aggressive mimicry. We computed an outer bound on *P. labiata*’s visual volume and found the periods at which the prey spider was in view and when it was not. Overall *P. labiata* could not see the prey for 9:57 (red portions of the speed vs. time profile); a total of 24% of the duration of the hunt, spanning 40% of the total path length. Scale bar is with respect to a feature near the bottom of the outermost capture spiral of the spider web (dark blue and green spheres in Supplementary Fig. 1).

### Aggressive Mimicry

Approximately one minute into the hunt, *P. labiata* crawled onto a strand of the web of its prey and appeared to engage in aggressive mimicry (Fig. 2; Video SM2a and SM2b). As shown in Fig. 2, *P. labiata* raised itself on the two rearmost pairs of legs, extending the two forward pairs of legs to reach onto the web silk (Video SM2a). *P. labiata* also intermittently moved its palps (Video SM2b). Although video resolution prevents us from confirming whether the palps contacted the web, *P. labiata* is known to use its palps in this way during aggressive mimicry^33^. *P. labiata* engaged in this behavior for almost 8 min before abandoning it and rerouting.

**Fig. 2:**
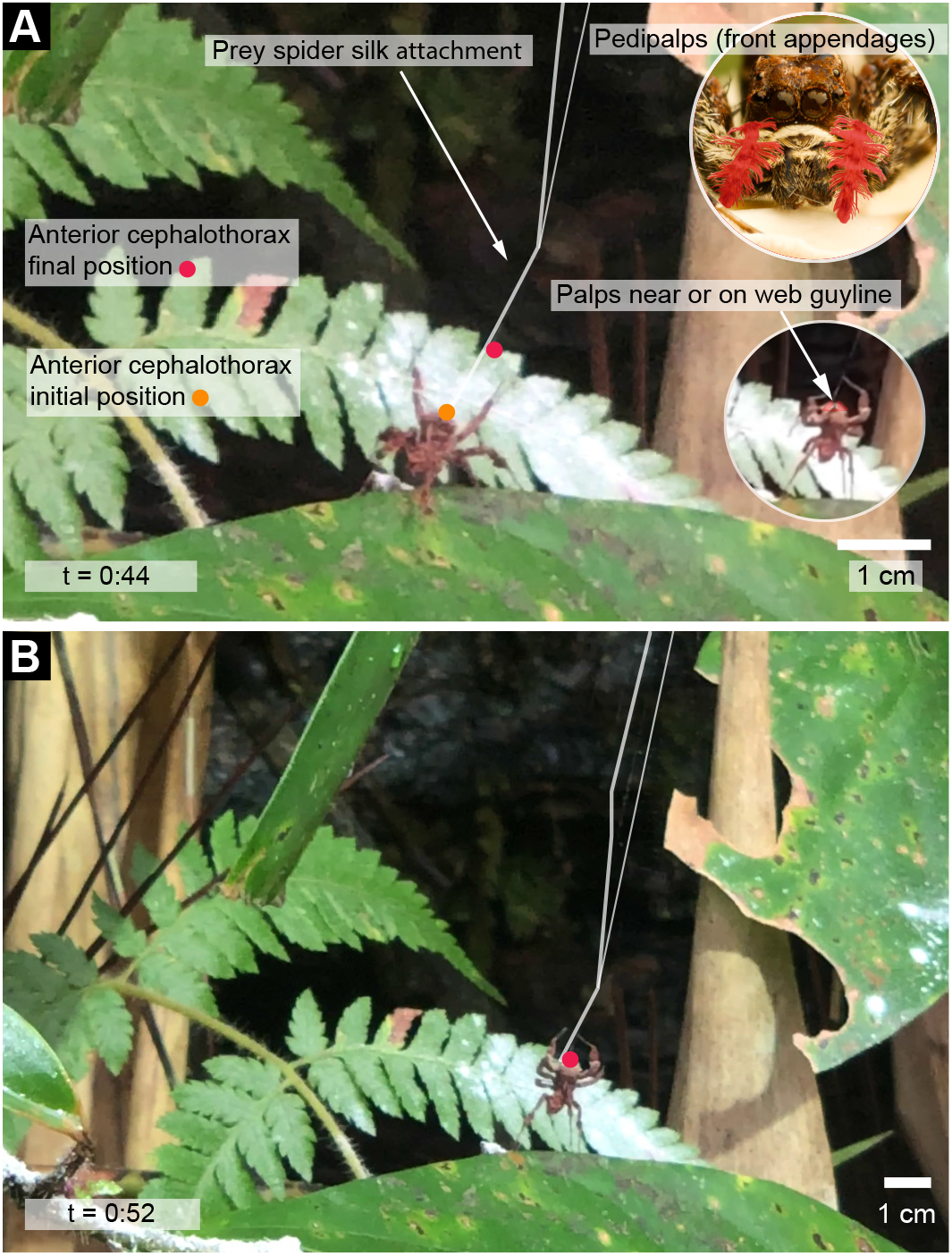
At about one minute into the hunt *Portia labiata* may have engaged in aggressive mimicry. **A**. *P. labiata* begins to crawl onto the web. The starting position of its cephalothorax is labeled in orange, and the final position is labeled in red. The palps in the inset images are artificially colored red to highlight them. **B**. *P. labiata* has positioned its first and second pairs of legs on the guyline. *P. labiata* appears to intermittently move its palps against the web (Video SM2a and SM2b). At 8:53 of SM1, *P. labiata* appears to give up on using aggressive mimicry and reroutes. Image of pedipalps courtesy of Marshall Patrick.

### Reroute

We computed the shortest path to the prey from *P. labiata*’s starting position after the end of the likely aggressive mimicry bout (Methods). Figure 3A shows this path going over a leaf with multiple guylines from the web connecting to it. Having abandoned the apparent attempt at aggressive mimicry, *P. labiata* first faced the leaf to its left (the shortest path to the prey), and then turned to scan and move to its right, along the longer path (Fig. 1, inset 2 & Video SM3 00:08–00:32). This path was almost double the shortest path it had originally oriented toward (212 mm versus 120 mm). Supplementary Table 5 outlines metrics between these two paths.

**Fig. 3:**
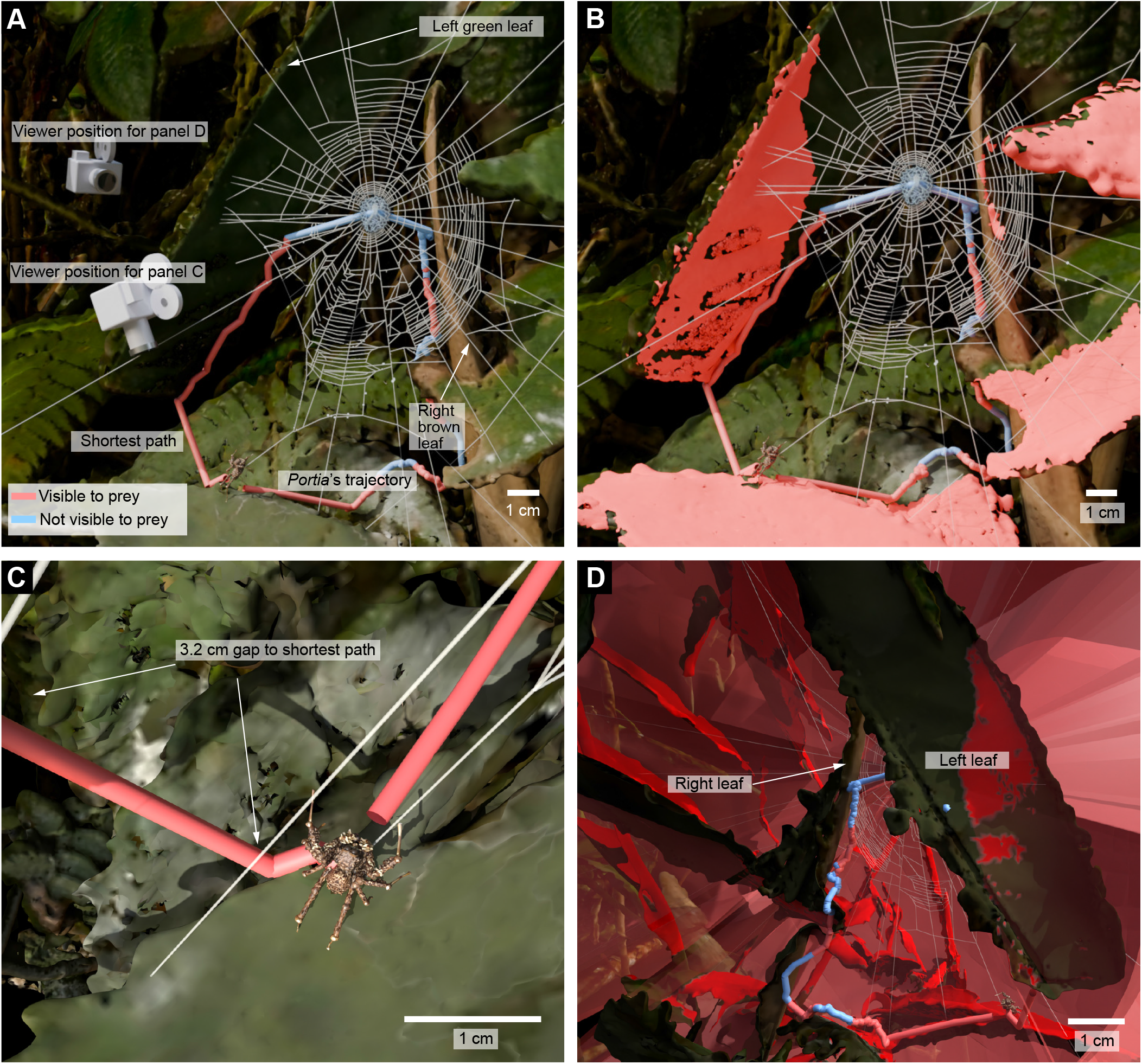
**Detour**: The detour after the end of the apparent aggressive mimicry attempt, along with one alternative path, the shortest possible to the prey. **A**. Shortest path compared to *P. labiata*’s trajectory (red is visible to prey on a generous estimate of its visual capacity, blue is not visible). Also shown are renderings of the camera view for panels **C** and **D. B**. Here we colored portions of the foliage consistent with an upper bound approximation of all locations where a 10 mm object (similar in size to *P. labiata*) could be potentially perceived. **C**. From the camera position shown in **A**, a close-up showing the 3.2 cm jump that would have been needed at the start of the shortest path. **D**. From the camera position shown in **A**, after rerouting, the blue portion of *P. labiata*’s trajectory was largely outside the estimated visual volume (red) unlike the red segments of the shortest path and the red first section of the hunt.

### Stalking

*Portia labiata* engaged in an almost imperceptibly slow stalk sequence, taking a total of 15 min to cover 4 cm before reaching its prey (Video SM4a; Fig. 4). *Portia labiata*’s movement was saltatory, where it paused for periods of roughly 3 min, before making a small, but abrupt movement forward (Video SM4b; Fig. 1). This slow stalking resembles the stalking sequences of mammals such as big cats, which are typically aware of their prey’s senses and attempt to minimize their own detectability^36, 37^.

**Fig. 4:**
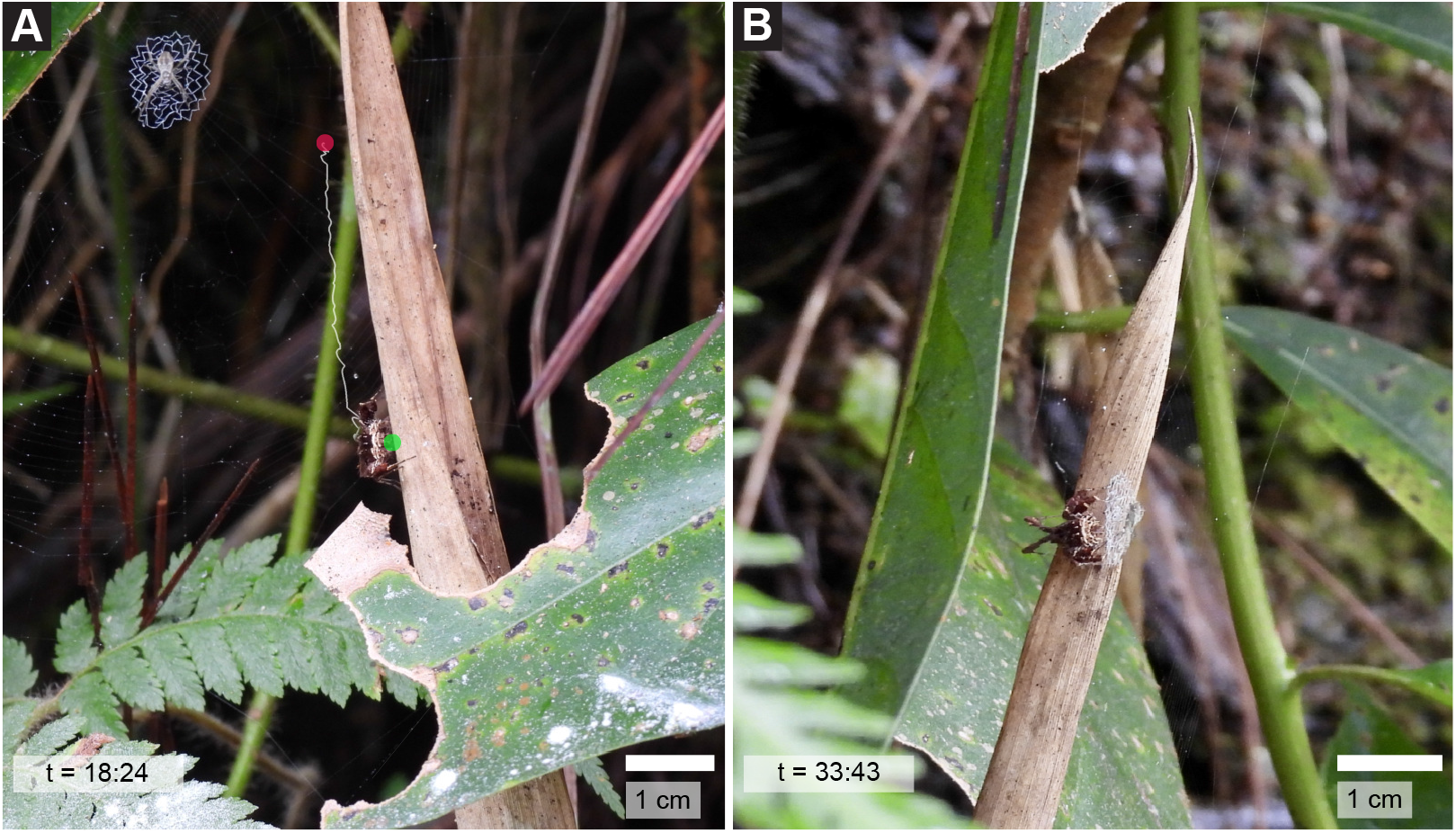
*Portia labiata* engages in a slow stalk, taking 15:03 to cover the 4 cm (from green to red dot) to reach the final position where it orients slowly for 05:05 before jumping at around 38:50 to attack the prey. **A**. *P. labiata* begins the stalk. The starting position of its cephalothorax is noted in green and its end position in red. **B**. At 33:47, *P. labiata* reaches the point from which it will attack a juvenile *Argiope versicolor*.

### Attack from under the discoid web decoration

Once *Portia labiata* reached the point from where it would jump at the prey, it spent 5 min slowly rolling and pitching its body to get into position for the final high speed lunge. We fully reconstructed *P. labiata*’s changing pose and leg position for this portion of the hunt (Supplementary 3D Animation SA1). *P. labiata* first moved into position by crab walking side to side (Video SM4b), then rolled and pitched its body to aim its attack (Video SM5a) and then raised its first and second pair of legs toward the prey in preparation for the attack lunge (Video SM5b). The lunge occurred at over 13,000 times the speed of the slow stalk.

As seen in Fig. 5 (and Videos SM5a, SM5b, and SM5c), *P. labiata* jumped from behind its prey’s web, landing on the ventral side of *A. versicolor* ‘s web discoid, and then pulled the prey spider through to the other side. To our knowledge this method of attack has not been previously reported. The purpose of the discoid is contested, although laboratory trials with an avian predator showed its presence made attacking from the ventral side less likely^4^.

**Fig. 5:**
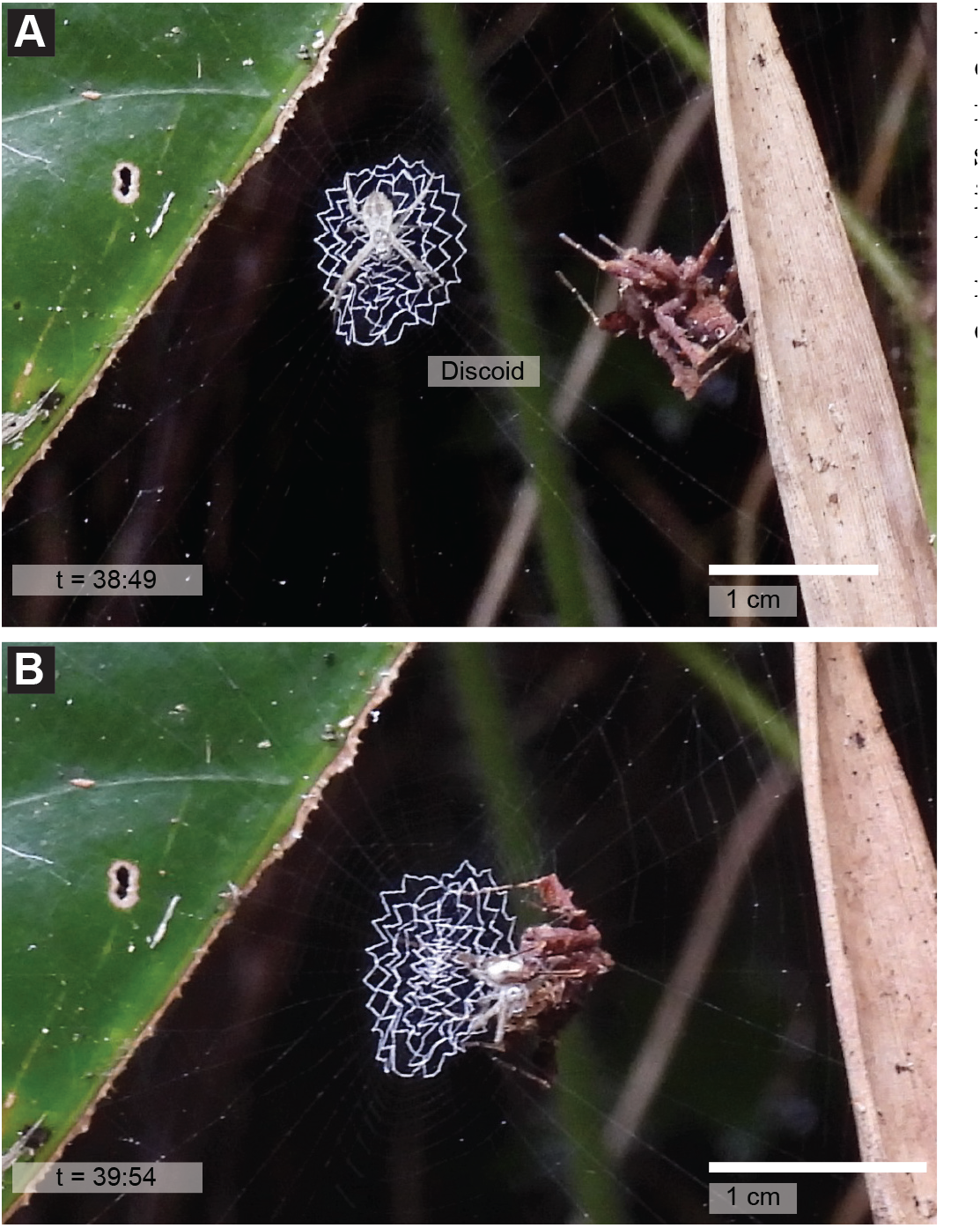
Novel attack from under the discoid web decoration: *Portia labiata* jumps from behind the web, landing on the opposite side of the discoid from the side the prey spider is positioned on, and pulls *Argiope versicolor* through its own web. **A**. *P. labiata* about to make its attack. **B**. *P. labiata* behind the web’s discoid just after attack.

## Discussion

### From potential aggressive mimicry to detouring

One possible signature of planning is the apparent strategy switch *P. labiata* makes early in the hunt. In the laboratory, *Portia* generates vibratory signals by trial and error when presented with unfamiliar prey (*Portia* species, including *P. labiata*, use trial and error for a range of strategies^28, 32, 38–40^), retaining those that elicit a response^28^. However, in this hunt, rather than continue aggressive mimicry, *P. labiata* switched tactics entirely after about 8 minutes. It then committed to a path about twice the length of the shortest available one and involving a period of nearly ten minutes where the prey was probably out of sight (Fig. 1, 09:20– 19:30 of SM1, likely out-of-view region colored blue in the velocity plot). We are not aware of such a tactic switch reported in a wild *P. labiata* hunt. While abandoning aggressive mimicry could be explained by a simple giving-up rule, where the spider gives up a tactic after it has failed for some threshold duration, this giving-up explanation does not fully explain the subsequent trajectory. In particular, *P. labiata*’s choice to avoid a substantially shorter path that may have generated more conspicuous visual and vibrational cues further suggests planning.

### Route choice in the predator-prey sensory landscape

We first consider visual and then mechanosensory factors. Starting with the prey, we estimate an upper bound on its visual range by using data from a spider whose visual acuity has been well characterized, a wandering spider *Cupiennius salei*, which likely has somewhat better vision than *A. versicolor* ^41, 42^ (Methods). *C. salei* relies primarily on mechanosensation rather than vision, broadly resembling *A. versicolor* in this respect^43–45^. While no measurements of acuity exist for *P. labiata*, there are for a closely-related species with similar morphology (*P. fimbriata*^46^). Using acuity measurements from *P. fimbriata*, and eye measurements of *P. labiata*^46^, we obtained an expected acuity of *≈* 10.5 cpd (Methods). See Supplementary Table 4 for outline of the vision models used in this study. We cannot resolve the exact position of *P. labiata*’s focal region during this hunt, since salticids direct gaze by moving the retinal tubes behind fixed corneal lenses^47^. Body orientation is nonetheless informative: on withdrawing from the web after the apparent failure of aggressive mimicry, *P. labiata* first oriented toward a left green vertical leaf along the shortest path, and it then turned approximately 90^*°*^away from it toward a right brown vertical leaf before starting the longer route (Video SM3 at 00:08– 00:32; Fig. 3A, left leaf and right leaf).

One possibility for why the longer route was taken is that *≈* 97% of the shorter route was in the perceptual envelope of the prey spider, while our estimates suggest less of the longer path would have been (blue segment, Fig. 3B,D). We note that this is based on the visual capacity of a spider likely to have better vision than *A. versicolor*.

The visual acuity estimate of *P. labiata* allows us to estimate that less of the longer path taken was visible to *P. labiata* at the rerouting decision point than the avoided shorter path (Methods), adding further challenge to this path.

Mechanosensory factors may have played a larger role in this route choice, however, given evidence that *P. labiata* avoids silk attachments (and the indirect evidence that it is sometimes successful in manipulating the prey based on vibrating one of those attachments). We know that *P. labiata* will skirt the edges of a web, and will very slowly and methodically cut silk attachments prior to an attack if they are in the way^33^. The left green leaf that *P. labiata* would have needed to crawl along for the shorter route is connected to the web by ten silk attachments (Fig. 3A). In addition, *P. labiata* would have had to make a 3.2 cm jump to the left leaf, double the 1.5 cm jump onto the right brown leaf that it made before it began its stalk (segment 2, high-lighted in purple, Fig. 1; Video SM1 at 12:15–13:40). Even for this 1.5 cm jump, *P. labiata* spent considerable time waving its first pair of legs toward the leaf and apparently reaching its body closer to the leaf, as if trying to crawl on to the leaf instead of jumping (Video SM1 at 13:15– 13:35; “hesitant jump,” Fig. 1). Although the 3.2 cm jump is well within *P. labiata*’s 8 cm forward-jumping capacity^33^, it may have caused movement of the leaf. Note that spiders, such as the prey spider of this hunt, have extremely sensitive mechanosensory systems^43, 44, 48^, making detection of vibrations on the leaf featuring many silk attachments quite possible.

Finally, after selection of the longer route, *P. labiata* approached its prey from behind the web, using a slow, saltatory stalking behavior (2.6 mm/min). This was the slowest *P. labiata* movement during the hunt. Both the slowness of the stalk and its location are notable. First, the speed means that any movements are unlikely to be detectable by either the prey’s mechanosensory system or motion-sensitive visual system. In addition, the path was well chosen, as this portion of the trajectory is mostly to the ventral side of the prey and thus is probably visually hidden from it, given that orb weaver eyes are directed dorsally and laterally^49^ and *A. versicolor* ‘s discoid shadows *P. labiata* from view.

### The adaptive significance of planning

In simulations testing the “Buena Vista” hypothesis mentioned earlier, in which visual range and environmental complexity were varied in a predator-prey scenario, planning had little value unless visual occlusion density was intermediate between a low density open plain and a high density jungle-like environment (where gaps between occlusions are similar to one body length)^16, 50^. Key to this intermediate level of occlusions was patchy clutter, where some positions offer concealment while others offer a view of the adversary. Two measures of the geometry of the foliage of this hunt indicate that the relevant spatial complexity is present in this hunt and four other nearby sites where we found *P. labiata*^16, 50^ (Methods; Supplementary Table 1, Fig. S2).

Planning is computationally costly^16^. Given a spider’s very small brain size, how could planning be implemented? We hypothesize that *P. labiata*’s extremely slow stop-and-go strategy may facilitate not only concealment^33, 51^, but allow opportunities to scan the environment and evaluate different outcomes. In *P. fimbriata*, the highest-acuity region of each principal-eye retina spans only about 0.6^*?*20^. Examining a larger field at that resolution involves scanning the retina across the scene and gathering information over seconds^24, 47^. Route evaluation could follow a similar principle: reusing limited neural resources to evaluate alternatives sequentially.

On this interpretation, the extended pauses and scanning bouts that characterize *P. labiata*’s hunts^6^ offer it moments to compute alternatives and tactics. Unlike the pauses of “vicarious trial and error” (VTE) in rodents, which are thought to reflect planning through an internal search rather than sensory sampling^1^, *Portia*’s scanning is overtly visual. It more closely resembles the gaze-based counterpart of VTE in visual animals—humans and other primates choosing between options look back and forth from one to another^1^. In some laboratory detour tasks mentioned earlier^6^, both routes were in view, one was impassable, and the spider picked the one without the gap, which it scanned the most. The hunt described here poses a different problem: both routes were passable, but the shorter route could have alerted the prey through its sensory capabilities. The longer route taken did not have silk attachments after the aggressive mimicry location, and was mostly outside the potential visual envelope of the prey, but at least a fifth of the route was also out of view of *P. labiata*— more so than the shortest route (Methods). Given that *P. labiata* could not see either of the paths in their entirety, its choice cannot have rested only on fixating the whole route it then followed. Looking may be part of how a visual animal evaluates a route, but for the parts it cannot see, the evaluation must draw on something other than the current view, such as an internal representation of the prey within a cognitive map of the hunting space.

### Conclusion

Planning has been studied most extensively in mammals, including classic observations of rats devising novel routes^52^. Despite decades of neurophysiology, there has yet to emerge a clear physiological criterion for when behavior counts as plan-driven as opposed to learned or innate routine-driven^13, 18^. Behavioral evidence alone is unlikely to establish that *P. labiata* can plan, but behavioral observations of its planning-like behavior are an important step. Similarly, work on problem-solving in bees does not show definitively^53^ that they are capable of flexible problem-solving, but it provides evidence bearing on that hypothesis^54, 55^.

For as complex a phenomenon as planning that spans many scientific domains, including ethology, psychology, computer science, robotics, and neuroscience, a pluralistic approach is needed^56^. To these dimensions it would be helpful to add considerations of why planning evolved, and to compare instantiations of it across animals with non-planning last common ancestors. Given the more than 600 million year gap between the last common ancestor of the arthropod and vertebrate lineages^9, 10, 57^, examining planning’s possible presence in invertebrates could help disentangle contingency from necessity. Is there a “physics of planning” that these animals can help illuminate, in the manner in which flapping flight in insects and vertebrates illuminates the physics-based necessity of flapping flight^58, 59^? Further work is needed, but *Portia*’s comparatively small nervous system may offer opportunities for rigorous studies of cognition, as *Drosophila* has in sensorimotor processing.

## Supplementary Animation and Movies

SA1: Interactive 3D animation (webGL) that permits moving through the time and volume of the hunt. https://maciverlab.github.io/portia-wild-hunt

For preprint/review purposes, given the size of the video files, we provide a link to a YouTube upload of each. The view may default to the low resolution version. To obtain the high resolution version, in the YouTube viewing interface, select the gear icon, select Quality, then select 1080p HD.

SM1: Synchronized videos of full hunt. https://youtu.be/y0AfnKGiONs

SM2a: *P. labiata* crawls onto the prey’s web at the beginning of the hunt, pauses, and holds it in tension. https://youtu.be/doiqBGyoOnk

SM2b: Potential aggressive mimicry. *Portia labiata* moves its palps apparently against the guyline. https://youtu.be/H16Q2mqNMXc

SM3: Reroute: *P. labiata* crawls off the web, orients itself toward the leaf that carries the shortest path, then turns 90^*°*^to take 11 a different route. https://youtu.be/ZPn8R1qksUQ

SM4a: Full stalk sequence. *Portia labiata* reaches a dry leaf and begins to walk up it to gain access to its prey. It engages in a slow predatory stalk. https://youtu.be/Mw1ZfCZUSUM

SM4b: Stalk sequence stabilized. Stalk sequence has been stabilized around the DLC tracked cephalothorax of *P. labiata*, sped up 50x. The saltatory approach becomes clear, as well as a short crab walk when positioning itself to jump on the prey. https://youtu.be/Uuy65WTHBYc

SM5a: *Portia labiata* rolls and pitches its body in preparation for attack. https://youtu.be/e27-w1iL-8U

SM5b: Final attack sequence. *Portia labiata* attacks from behind the web, lands on the ventral side of the prey’s discoid, and pulls the prey through from the dorsal side of the web to the ventral side. https://youtu.be/Qc6MMPrYCk8

SM5c: Final attack sequence reconstructed. Rendered at 24x speed in Blender 5.1. Pose reconstructed using full pose tracking via DLC and position of web and spider found using Structure from Motion. https://youtu.be/dLQBejDRgyI

SM6: Stalk video sequence aligned to the laser scan, displayed 10 frames per second. Used to confirm alignment accuracy. https://youtu.be/ncIRF3Kn_DY

## Acknowledgements

This work was funded by a grant from the Human Frontiers Scientific Program, RGP017/2024, to CL, XN, DL, and MM. We thank Shonan 3D Solutions Pte Ltd, Singapore, for assistance with 3D scanning of the foliage and spider. We thank Prof. Emma Alexander, Department of Computer Science, Northwestern University, for guidance on computer vision methods. We also thank Eugene Lin, Marshall Patrick, and Yingna Zhou who performed the manual orientation (positioning *P. labiata* in space, and in yaw, pitch, roll for 286 frames). Further thanks to Marshall Patrick for supplying the *P. labiata* picture used to illustrate the pedipalps in Fig. 2. Finally we thank Alex Lai, Lucas Lippeatt, and Izzy Bider for correcting DLC estimates in the final attack reconstruction sequence.

## Resource availability Lead contact

Further information and requests for resources should be directed to and will be fulfilled by the lead contact, Dr. Malcolm MacIver

## Declaration of generative AI and AI-assisted technologies

During the preparation of this work, the authors used ChatGPT and Claude for generating alternative phrasings of sentences and shortening text. We also used Claude for code assistance. After using this tool, the authors reviewed and edited the content as needed and take full responsibility for the content of the publication.

## Footnotes

1 This and subsequent links to Supplementary Videos are hyperlinks within the PDF to YouTube uploads for reviewer convenience given the size of the videos.

## References

1. Redish, A.D. (2016). Vicarious trial and error. Nature Reviews Neuroscience 17, 147–159. 10.1038/nrn.2015.30.

2. Güntürkün, O. and Bugnyar, T. (2016). Cognition without cortex. Trends in Cognitive Sciences 20, 291–303. 10.1016/j.tics.2016.02.001.

3. Raby, C.R., Alexis, D.M., Dickinson, A., and Clayton, N.S. (2007). Planning for the future by western scrub-jays. Nature 445, 919–921. 10.1038/nature05575.

4. Wang, B., Yu, L., Ma, N., Zhang, Z., Liu, Q., Fan, W., Rong, Y., Zhang, S., and Li, D. (2021). Discoid decorations function to shield juvenile Argiope spiders from avian predator attacks. Behavioral Ecology 32, 1230–1239. 10.1093/beheco/arab089.

5. Tarsitano, M.S. and Jackson, R.R. (1997). Araneophagic jumping spiders discriminate between detour routes that do and do not lead to prey. Animal Behaviour 53, 257–266. 10.1006/anbe.1996.0372.

6. Tarsitano, M.S. and Andrew, R. (1999). Scanning and route selection in the jumping spider Portia labiata. Animal Behaviour 58, 255–265. 10.1006/anbe.1999.1138.

7. Cross, F.R., Carvell, G.E., Jackson, R.R., and Grace, R.C. (2020). Arthro-pod intelligence? The case for Portia. Frontiers in Psychology 11, 568049. 10.3389/fpsyg.2020.568049.

8. Chittka, L. and Niven, J. (2009). Are bigger brains better? Current Biology 19, R995–R1008.

9. Erwin, D.H., Laflamme, M., Tweedt, S.M., Sperling, E.A., Pisani, D., and Peterson, K.J. (2011). The Cambrian Conundrum: Early Divergence and Later Ecological Success in the Early History of Animals. Science 334, 1091–1097. 10.1126/science.1206375.

10. dos Reis, M., Thawornwattana, Y., Angelis, K., Telford, M.J., Donoghue, P.C.J., and Yang, Z. (2015). Uncertainty in the Timing of Origin of Animals and the Limits of Precision in Molecular Timescales. Current Biology 25, 2939–2950. 10.1016/j.cub.2015.09.066.

11. Evans, S.D., Hughes, I.V., Gehling, J.G., and Droser, M.L. (2020). Discovery of the oldest bilaterian from the Ediacaran of South Australia. Proceedings of the National Academy of Sciences 117, 7845–7850. 10.1073/pnas.2001045117.

12. Nilsson, D.E. (2013). Eye evolution and its functional basis. Visual Neuroscience 30, 5–20. 10.1017/S0952523813000035.

13. Mattar, M.G. and Lengyel, M. (2022). Planning in the brain. Neuron 110, 914–934. 10.1016/j.neuron.2021.12.018.

14. MacIver, M.A. (2009). Neuroethology: From Morphological Computation to Planning. In The Cambridge Handbook of Situated Cognition, P. Robbins and M. Aydede, eds. (New York, NY: Cambridge University Press), pp. 480–504.

15. MacIver, M.A., Schmitz, L., Mugan, U., Murphey, T.D., and Mobley, C.D. (2017). Massive increase in visual range preceded the origin of terrestrial vertebrates. Proceedings of the National Academy of Sciences 114, E2375–E2384. 10.1073/pnas.1615563114.

16. Mugan, U. and MacIver, M.A. (2020). Spatial planning with long visual range benefits escape from visual predators in complex naturalistic environments. Nature Communications 11, 3057. 10.1038/s41467-020-16102-1.

17. MacIver, M.A. and Finlay, B.L. (2022). The neuroecology of the water-to-land transition and the evolution of the vertebrate brain. Philosophical Transactions of the Royal Society B: Biological Sciences 377, 20200523. 10.1098/rstb.2020.0523.

18. Hunt, L.T., Daw, N.D., Kaanders, P., MacIver, M.A., Mugan, U., Procyk, E., Redish, A.D., Russo, E., Scholl, J., Stachenfeld, K., et al. (2021). Formalizing planning and information search in naturalistic decision-making. Nature Neuroscience 24, 1051–1064. 10.1038/s41593-021-00866-w.

19. Fleming, S.M. and Michel, M. (2025). Sensory Horizons and the Functions of Conscious Vision. Behavioral and Brain Sciences 49, e162. 10.1017/S0140525X25000068.

20. Williams, D.S. and McIntyre, P. (1980). The principal eyes of a jumping spider have a telephoto component. Nature 288, 578–580. 10.1038/288578a0.

21. Caves, E.M., Brandley, N.C., and Johnsen, S. (2018). Visual Acuity and the Evolution of Signals. Trends in Ecology & Evolution 33, 358–372. 10.1016/j.tree.2018.03.001.

22. Dunning, Jr., J.B. (2008). CRC Hand-book of Avian Body Masses. 2nd edition (Boca Raton, FL: CRC Press).

23. Sherk, T.E. (1978). Development of the compound eyes of dragonflies (Odonata). III. Adult compound eyes. Journal of Experimental Zoology 203, 61–79. 10.1002/jez.1402030107.

24. Harland, D.P., Li, D., and Jackson, R.R. (2012). How Jumping Spiders See the World. In How Animals See the World: Comparative Behavior, Biology, and Evolution of Vision, O.F. Lazareva, T. Shimizu, and E.A. Wasserman, eds. (Oxford: Oxford University Press), pp. 133–163.

25. Land, M.F. and Barth, F.G. (1992). The quality of vision in the ctenid spider Cupiennius salei. Journal of Experimental Biology 164, 227–242. 10.1242/jeb.164.1.227.

26. Tolman, E.C. (1948). Cognitive Maps in Rats and Men. Psychological Review 55, 189–208. 10.1037/h0061626.

27. Jackson, R.R. and Wilcox, R.S. (1993). Observations in nature of detouring behaviour by Portia fimbriata, a web-invading aggressive mimic jumping spider from Queensland. Journal of Zoology 230, 135–139.

28. Jackson, R.R. and Wilcox, R.S. (1993). Spider flexibly chooses aggressive mimicry signals for different prey by trial and error. Behaviour 127, 21–36.

29. Jackson, R.R. and Wilcox, R.S. (1990). Aggressive mimicry, prey-specific predatory behaviour and predator-recognition in the predator–prey interactions of Portia fimbriata and Euryattus sp., jumping spiders from Queensland. Behavioral Ecology and Sociobiology 26, 111–119.

30. Jackson, R.R. and Pollard, S.D. (1996). Predatory behavior of jumping spiders. Annual Review of Entomology 41, 287–308.

31. Tarsitano, M., Jackson, R.R., and Kirchner, W.H. (2000). Signals and Signal Choices made by the Araneophagic Jumping Spider Portia fimbriata while Hunting the Orb-Weaving Web Spiders Zygiella x-notata and Zosis geniculatus. Ethology 106, 595–615. 10.1046/j.1439-0310.2000.00570.x.

32. Jackson, R.R. and Nelson, X.J. (2011). Reliance on trial and error signal derivation by Portia africana, an ara-neophagic jumping spider from East Africa. Journal of Ethology 29, 301–307. 10.1007/s10164-010-0258-5.

33. Jackson, R.R. and Hallas, S.E.A. (1986). Comparative biology of Portia africana, P. albimana, P. fimbriata, P. labiata, and P. shultzi , araneophagic, web-building jumping spiders (Araneae: Salticidae): Utilisation of webs, predatory versatility, and intraspecific interactions. New Zealand Journal of Zoology 13, 423–489. 10.1080/03014223.1986.10422978.

34. Jackson, R.R., Li, D., Fijn, N., and Barrion, A. (1998). Predator–prey interactions between aggressive-mimic jumping spiders (Salticidae) and ara-neophagic spitting spiders (Scytodidae) from the Philippines. Journal of Insect Behavior 11, 319–342. 10.1023/A:1020946529246.

35. Jackson, R.R., Pollard, S.D., Li, D., and Fijn, N. (2002). Interpopulation Variation in the Risk-Related Decisions of Portia labiata, an Araneophagic Jumping Spider (Araneae, Salticidae), during Predatory Sequences with Spitting Spiders. Animal Cognition 5, 215–223. 10.1007/s10071-002-0150-y.

36. Latest Sightings (2021). Buck Doesn’t See the Leopard Behind This Tree … YouTube video. URL https://www.youtube.com/watch?v=zReSMP1pM5M.

37. Elliott, J.P., Cowan, I.M., and Holling, C.S. (1977). Prey capture by the African lion. Canadian Journal of Zoology 55, 1811–1828. 10.1139/z77-235.

38. Cross, F.R. and Jackson, R.R. (2015). Solving a novel confinement problem by spartaeine salticids that are predisposed to solve problems in the context of predation. Animal Cognition 18, 509–515.

39. Jackson, R.R., Carter, C.M., and Tar-sitano, M.S. (2001). Trial-and-error solving of a confinement problem by a jumping spider, Portia fimbriata. Behaviour 138, 1215–1234. 10.1163/15685390152822184.

40. Jackson, R.R., Cross, F.R., and Carter, C.M. (2006). Geographic variation in a spider’s ability to solve a confinement problem by trial and error. International Journal of Comparative Psychology 19, 282–296. 10.46867/ijcp.2006.19.03.06.

41. Morehouse, N.I., Buschbeck, E.K., Zurek, D.B., Steck, M., and Porter, M.L. (2017). Molecular evolution of spider vision: new opportunities, familiar players. The Biological Bulletin 233, 21–38. 10.1086/693977.

42. Land, M.F. (1985). The morphology and optics of spider eyes. In Neurobiology of Arachnids, F.G. Barth, ed. (Berlin, Heidelberg: Springer), pp. 53–78.

43. Barth, F.G. (2002). A Spider’s World: Senses and Behavior (Berlin, Heidelberg: Springer).

44. Barth, F.G. (2004). Spider mechanore-ceptors. Current Opinion in Neurobiology 14, 415–422. 10.1016/j.conb.2004.07.005.

45. Hergenröder, R. and Barth, F.G. (1983). Vibratory signals and spider behavior: how do the sensory inputs from the eight legs interact in orientation? Journal of Comparative Physiology A 152, 361–371. 10.1007/BF00606241.

46. Wanless, F.R. (1978). A revision of the spider genus Portia (Araneae: Salticidae). Bulletin of the British Museum (Natural History), Zoology 34, 83–124.

47. Land, M.F. (1969). Movements of the retinae of jumping spiders (Salticidae: Dendryphantinae) in response to visual stimuli. Journal of Experimental Biology 51, 471–493. 10.1242/jeb.51.2.471.

48. Landolfa, M.A. and Barth, F.G. (1996). Vibrations in the orb web of the spider Nephila clavipes: cues for discrimination and orientation. Journal of Comparative Physiology A 179, 493–508. 10.1007/BF00192316.

49. Eberhard, W.G. (2017). How orb-weavers find and grasp silk lines. Journal of Arachnology 45, 145–151. 10.1636/JoA-S-16-057.1.

50. Lai, A.T., Espinosa, G., Wink, G.E., Angeloni, C.F., Dombeck, D.A., and MacIver, M.A. (2024). A robot-rodent interaction arena with adjustable spatial complexity for ethologically relevant behavioral studies. Cell reports (Cambridge) 43, 113671–113671.

51. Wilcox, R.S., Jackson, R.R., and Gentile, K. (1996). Spiderweb smoke-screens: spider trickster uses back-ground noise to mask stalking movements. Animal Behaviour 51, 313–326. 10.1006/anbe.1996.0031.

52. Tolman, E.C., Ritchie, B.F., and Kalish, D. (1946). Studies in Spatial Learning. I. Orientation and the Short-Cut. Journal of Experimental Psychology 36, 13–24. 10.1037/h0053944.

53. Alem, S., Perry, C.J., Zhu, X., Loukola, O.J., Ingraham, T., Søvik, E., and Chit-tka, L. (2016). Associative mechanisms allow for social learning and cultural transmission of string pulling in an insect. PLOS Biology 14, e1002564. 10.1371/journal.pbio.1002564.

54. Loukola, O.J., Solvi, C., Coscos, L., and Chittka, L. (2017). Bumblebees show cognitive flexibility by improving on an observed complex behavior. Science 355, 833–836.

55. Bhambore, A.A., Akmeşe, E.N., Häkki-nen, E., Jussila, M.K., Kantola, J.H., and Loukola, O.J. (2026). Spontaneous problem-solving in bumble bees. Science 392, 1046–1049. 10.1126/science.ady1618.

56. Krakauer, J.W., Ghazanfar, A.A., Gomez-Marin, A., MacIver, M.A., and Poeppel, D. (2017). Neuroscience Needs Behavior: Correcting a Reductionist Bias. Neuron (Cambridge, Mass.) 93, 480–490.

57. Kumar, S., Stecher, G., Suleski, M., and Hedges, S.B. (2017). TimeTree: A Resource for Timelines, Timetrees, and Divergence Times. Molecular Biology and Evolution 34, 1812–1819. 10.1093/molbev/msx116.

58. Lentink, D. and Dickinson, M.H. (2009). Biofluiddynamic scaling of flapping, spinning and translating fins and wings. Journal of Experimental Biology 212, 2691–2704. 10.1242/jeb.022251.

59. Chin, D.D. and Lentink, D. (2016). Flapping wing aerodynamics: from insects to vertebrates. Journal of Experimental Biology 219, 920–932. 10.1242/jeb.042317.

60. Schönberger, J.L. and Frahm, J.M. (2016). Structure-from-Motion Revisited. In IEEE Conference on Computer Vision and Pattern Recognition (CVPR). pp. 4104–4113.

61. Pan, L., Baráth, D., Pollefeys, M., and Schönberger, J.L. (2024). Global Structure-from-Motion Revisited. In European Conference on Computer Vision (ECCV).

62. Fischler, M.A. and Bolles, R.C. (1981). Random sample consensus: a paradigm for model fitting with applications to image analysis and automated cartography. Communications of the ACM 24, 381–395. 10.1145/358669.358692.

63. Umeyama, S. (1991). Least-squares estimation of transformation parameters between two point patterns. IEEE Transactions on Pattern Analysis and Machine Intelligence 13, 376–380. 10.1109/34.88573.

64. Besl, P.J. and McKay, N.D. (1992). A method for registration of 3-D shapes. IEEE Transactions on Pattern Analysis and Machine Intelligence 14, 239–256. 10.1109/34.121791.

65. Zhou, Q.Y., Park, J., and Koltun, V. (2018). Open3D: A Modern Library for 3D Data Processing.

66. Mathis, A., Mamidanna, P., Cury, K.M., Abe, T., Murthy, V.N., Mathis, M.W., and Bethge, M. (2018). DeepLabCut: markerless pose estimation of user-defined body parts with deep learning. Nature Neuroscience 21, 1281–1289. 10.1038/s41593-018-0209-y.

67. Nath, T., Mathis, A., Chen, A.C., Patel, A., Bethge, M., and Mathis, M.W. (2019). Using DeepLabCut for 3D markerless pose estimation across species and behaviors. Nature Protocols 14, 2152–2176. 10.1038/s41596-019-0176-6.

68. Virtanen, P., Gommers, R., Oliphant, T.E., Haberland, M., Reddy, T., Cour-napeau, D., Burovski, E., Peterson, P., Weckesser, W., Bright, J., et al. (2020). SciPy 1.0: fundamental algorithms for scientific computing in Python. Nature Methods 17, 261–272. 10.1038/s41592-019-0686-2.

69. Caves, E.M. and Johnsen, S. (2018). AcuityView: An R package for portraying the effects of visual acuity on scenes observed by an animal. Methods in Ecology and Evolution 9, 793–797. 10.1111/2041-210X.12911.

70. Hart, P.E., Nilsson, N.J., and Raphael, B. (1968). A Formal Basis for the Heuristic Determination of Minimum Cost Paths. IEEE Transactions on Systems Science and Cybernetics 4, 100–107. 10.1109/TSSC.1968.300136.

71. Mandelbrot, B.B. (1982). The Fractal Geometry of Nature (San Francisco: W. H. Freeman).

72. Plotnick, R.E., Gardner, R.H., and O’Neill, R.V. (1993). Lacunarity indices as measures of landscape texture. Landscape Ecology 8, 201–211. 10.1007/BF00125351.

73. Sebok, D., Vasarhelyi, L., Szenti, I., Vajtai, R., Konya, Z., and Kukovecz, A. (2021). Fast and accurate lacunarity calculation for large 3D micro-CT datasets. ACTA MATERIALIA 214. 10.1016/j.actamat.2021.116970.

74. Plotnick, R.E., Gardner, R.H., Hargrove, W.W., Prestegaard, K., and Perlmutter, M. (1996). Lacunarity analysis: a general technique for the analysis of spatial patterns 53, 5461.

75. Hoechstetter, S., Walz, U., and Thinh, N.X. (2011). Adapting lacunarity techniques for gradient-based analyses of landscape surfaces 8, 229–238.

76. DeTone, D., Malisiewicz, T., and Rabinovich, A. (2018). SuperPoint: Self-Supervised Interest Point Detection and Description. In Proceedings of the IEEE/CVF Conference on Computer Vision and Pattern Recognition Work-shops (CVPRW). pp. 224–236.

77. Lindenberger, P., Sarlin, P.E., and Pollefeys, M. (2023). LightGlue: Local Feature Matching at Light Speed. In Proceedings of the IEEE/CVF International Conference on Computer Vision (ICCV). pp. 17627–17638.

78. Uehara, A., Toh, Y., and Tateda, H. (1977). Fine structure of the eyes of orb-weavers, Argiope amoena L. Koch (Araneae: Argiopidae). 1. The anteromedial eyes. Cell and Tissue Research 182, 81–91. 10.1007/BF00222056.

79. Uehara, A., Toh, Y., and Tateda, H. (1978). Fine structure of the eyes of orb-weavers, Argiope amoena L. Koch (Aranea: Argiopidae). 2. The anterolateral, posteromedial and posterolateral eyes. Cell and Tissue Research 186, 435–452.

