## Supplementary methods and tables for "Can a Tiny Brain Plan? Reconstructing a Predatory Detour in the Jumping Spider *Portia labiata*"

### Methods and Supplementary Material

### Methods

The coordinates of the hunt location were 4.4757° N, 101.3831° E, 1,464 m elevation. The *P. labiata* was 6.3 mm in length (measured from the tip of the abdomen to the front of its cephalothorax). Its mass was not measured, but a set of *P. labiata* that were between 6–7 mm collected for another study weighed 47±2.5 mg (N=20, D. Li unpubl. data). The data from the hunt included a laser scan of the hunt's surrounding foliage and two videos of the spider's movements. The laser scanner was a handheld Artec Leo 3D scanner (Artec Europe S.à. r.l. Senningerberg, Luxembourg), with an accuracy of 0.1 mm and 3D resolution of 0.2 mm. The scanning mode was set to Geometry and texture using HD mode, with high resolution enabled. The foliage scan was carried out at 40 frames $s^{-1}$ and resulted in a mesh with 9 million polygons. Processing was performed in Artec Studio Professional (v 18.1.4.9, Artec Europe). After post-processing, including a cleanup process, and mesh generation (fusion resolution 0.5–0.7 mm), the foliage model was 4,493,317 vertices or 2 million polygons. A female specimen of *P. labiata* (not the one from the hunt) was digitally scanned using photogrammetry. Photographs were taken using the rear camera of a POCO X6 Pro smartphone (Xiaomi Communications Co., Ltd., Beijing, China), with the flash disabled. To minimize shadows and provide uniform illumination, the spider was photographed inside a PULUZ photo light box (Shenzhen PULUZ Technology Limited, Shenzhen, China). Images were processed with the photogrammetry capabilities within Artec Studio (v 19 Beta).

During the hunt, two members of the team used iPhones to record video. One video was recorded at 1080 × 1920 pixels and 59.97 frames $s^{-1}$ using an iPhone 12 Pro (Apple Inc., Cupertino, California, USA); the other was recorded at 1920 × 1080 pixels and 30.00 frames $s^{-1}$ using an iPhone X (Apple Inc.). Times are given as MM:SS (minutes:seconds) from the start of the associated video (including opening title card). Another member took a total of 18 photos on a Nikon COOLPIX P950; these were used for figures, as well as web reconstruction.

We reconstructed *P. labiata*'s trajectory through a combination of manual and automatic methods, and rendered the result in Blender (version 5.1; Blender Foundation, Amsterdam, The Netherlands) using the Cycles path tracing engine. We reconstructed the hunt scene from both videos with the open-source Structure from Motion (SfM) packages COLMAP[60] and GLOMAP[61]. The iPhone 12 Pro video was downsampled to 30 frames $s^{-1}$ (Premiere Pro with default settings). We extracted frames from each video at 1 frame $s^{-1}$. Afterwards the iPhone 12 Pro's frames were reconstructed with COLMAP's incremental mapper. The much longer iPhone X sequence fragmented under incremental mapping, so we reconstructed it with GLOMAP's global mapper instead. Each sparse model gave the cameras' positions over time along with the prey's web and position, and dense reconstruction then added detail to both point clouds. To recover a camera pose at regular intervals, we sampled each video at 5 Hz and registered these frames into its model by absolute pose estimation. Finally, we registered the two reconstructions against each other to merge them into a single point cloud. The photogrammetry model of *P. labiata* was scaled to match hunt's *P. labiata* in the SfM point cloud after alignment to the laser scan. Software versions and all reconstruction settings are given in Supplementary Table 2.

We fit the web plane with RANSAC (Random Sample Consensus; custom Python 3.13 script)[62] on a manually selected region of the point cloud belonging to the web, and used RANSAC again to recover the web strands present in the point cloud. Strands that SfM did not capture were traced in frames from the iPhone 12 Pro video and the Nikon COOLPIX P950 and projected onto the web plane. To make the silk visible for tracing, each frame was enhanced with a morphological (white top-hat) filter (OpenCV 4.12) that keeps bright structures narrower than the threads and stabilimentum and suppresses broader ones such as leaves and the background. We then contrast-stretched the result and overlaid it on a darkened copy of the frame. To find where the web anchored to the environment, we took the first intersection with the foliage along each radial's direction.

SfM is scale-free, so to recover metric scale we aligned the SfM point cloud to the laser scan. We first solved for the transformation mapping six corresponding features between the two (Procrustes analysis[63]), then adjusted the orientation by hand and refined the alignment with iterative closest point (ICP)[64], implemented in the open-source Python package Open3D (0.19.0)[65]. The median distance from each laser-scan point to the nearest point of the aligned SfM point cloud was 0.41 mm (interquartile range 0.12–1.22 mm; mean $0.88 \pm 1.13$ mm; Supplementary Fig. 1B). We also confirmed the alignment by rendering the cameras' frames over the laser scan (Video SM6; Supplementary Fig. 1A). At this scale, the prey *A. versicolor* measured $\approx$6 mm from the tips of its front to rear legs, and its discoid $\approx$9 mm in diameter (Blender measuring tool). Software settings for these steps are given in Supplementary Table 2.

To track *P. labiata*'s trajectory on the 3D reconstruction of the foliage, we combined DeepLabCut (DLC 2.3)[66, 67] with manual estimation. DLC tracked the spider's cephalothorax and abdomen throughout the hunt, and in the final capture also three points on each leg (coxa, tibia and tarsus). From these tracked points and the camera poses and intrinsics, we solved for *P. labiata*'s position as the least-squares convergence point of the viewing rays from adjacent frames. During the stalk, which was filmed by both cameras, we instead took the intersection of the two cameras' simultaneous viewing rays (custom Python 3.13 script). For the full-pose reconstruction of the final capture, we solved for the body's position in the same way, then used inverse kinematics to pose the legs touching the dry leaf and forward kinematics for legs in the air, successively locating each point of the tracked leg chain (coxa, tibia and tarsus). Four annotators (two experts on *Portia* species and two non-experts) then reviewed 286 frames across the hunt, correcting the estimated positions and estimating the spider's orientation, which DLC did not give robustly. They also marked where the automatic method failed, such as when the spider was behind foliage or moving too fast for rays from adjacent frames to converge; 157 frames were estimated by annotation alone. For the final attack, the four annotators oriented *P. labiata* over 16 frames (Supplementary Fig. 1).

We smoothed *P. labiata*'s velocity with a non-causal Savitzky–Golay filter (SciPy 1.17)[68]. Because *P. labiata* hunts very slowly, this acted as a low-pass filter, removing high-frequency noise from tracking and alignment while keeping the spider's movement, but it also removed the brief jump and final attack. For the velocity plot in Fig. 1, we therefore estimated the speeds of the hesitant jump (450 mm $s^{-1}$) and the final attack (600 mm $s^{-1}$) from the spider's displacement across three consecutive frames. Tracking and smoothing settings are given in Supplementary Table 3.

We did not track *P. labiata*'s legs outside of the final attack. For the web animation its gait was generated procedurally with inverse kinematics (IK) using a custom script in Blender. The body follows the reconstructed 3D trajectory, and the script animates only the eight foot targets; the IK solver then sets the leg joint angles given these targets.

Principal-eye proportions relative to the anterior eye row are nearly identical in females of *P. fimbriata* and *P. labiata*[46]. Scaling from the 0.81 mm corneal lens of the *P. fimbriata* eye whose acuity was measured at 12.5 cpd[20] to the ≈0.68 mm lens estimated for a *P. labiata* female of similar size to ours (6.5 vs 6.3 mm[46]), and assuming similar receptor spacing, gives an expected acuity of ≈10.5 cpd for *P. labiata*.

Using our 10.5 cpd acuity estimate, *P. labiata* can resolve the prey (measured as approximately 6 mm) from 3.6 m away, and is at a maximum distance of 11 cm from the prey. However, occlusions intermittently block visual access. Given the trajectory and orientation of the spider, we therefore computed the positions in which *P. labiata* loses sight of the prey due to occlusions. High acuity regions of the anterior-median eyes cover 0.6°[20]. Salticidae can move their retinal tube 50° horizontally, and 35° vertically (measured in *Metaphidippus*[47]). Furthermore *P. labiata* can roll and pitch its cephalothorax about 30° (X. Nelson, personal observations). We cannot be sure of *P. labiata*'s focal point, so we checked for line of sight within this wider potential view for all positions along the trajectory. Line of sight was computed by tracing rays within this field of view and checking whether these rays intersected with the prey spider. If there was an intersection, we then checked if there was a foliage intersection before the prey spider. We also computed how much of each candidate route (the shortest path and the path *P. labiata* actually took) was visible to *P. labiata* before it rerouted. Each route was represented as a tube of 10 mm diameter (an upper bound on *P. labiata*'s body length) around its centreline. Using the line-of-sight computation described above, we determined which parts of each tube were visible from *P. labiata*'s position every 500 frames (≈ 32 s intervals) from the start of the hunt to the rerouting point (frames 0–8254, including the final frame). A part of a route was counted as seen if it was visible in at least one of these frames, and we report the seen fraction of each route's length.

To our knowledge, no visual acuity measurements exist for any member of the prey spider's family, Araneidae. Because web-building spiders have reduced eyes relative to wandering spiders[41, 42], we used the anatomically derived acuity of the wandering spider *Cupiennius salei* as an upper bound on our juvenile *A. versicolor*. For example, *C. salei*'s anterior lateral eyes are the lowest resolution, with inter-receptor angles of $\Delta\phi = 3.6°$ along the receptor rows[25], giving a Nyquist limit of $\nu = 1/(2\Delta\phi) = 0.139\,\text{cpd}$. For a 10 mm object (an upper bound on the size of *P. labiata*, which ranges from 6 mm to 10 mm), we calculated visual range as follows. Assuming unity contrast, an object of size $s$ is resolvable by an eye of acuity $\nu$ (in cycles per degree) only if it subtends at least one full cycle, $\theta = 1/\nu$[21]. The corresponding maximum resolvable distance is[69]

$$d = \frac{s}{2\tan\left(\frac{\theta}{2}\right)} = \frac{s}{2\tan\left(\frac{1}{2\nu}\right)}, \quad (1)$$

where the argument of the tangent is in degrees. With *Cupiennius salei* as an upper bound, $s = 10\,\text{mm}$ and its anterior-lateral eye acuity measured as $\nu = 0.139\,\text{cpd}$, this gives $\theta = 7.2°$ and $d = 79.5\,\text{mm}$.

We computed the maximum resolvable distance for the four pairs of eyes which give distances of 9.9 cm (anterior median), 7.9 cm (anterior lateral), 31.8 cm (posterior median), and 28.6 cm (posterior lateral). We used Land's and Barth's map for the angles of the field of view of these eyes[25] and deprojected these regions using Hammer-Aitoff projection[25] to isolate the field of views of these eyes in 3-dimensions. The position of *A. versicolor*'s eyes was taken as its head in the reconstruction. We allowed *A. versicolor* to see 40 degrees beneath the web plane based on the angle below the horizontal for which *C. salei* is able to see[25]. We blocked lines of sight that intersected *A. versicolor*'s discoid. We calculated the intersection of these volumes with the surrounding geometry, as well as *P. labiata*'s path. A point was classified as visible to the prey if it lay inside the field of view of at least one eye, within that eye's maximum resolvable distance (for the 10 mm upper bound of *Portia* size), and had an unobstructed line of sight to the prey's eyes, tested by ray casting against the laser scan of the foliage as well as the prey's discoid.

To solve for the shortest route available to *P. labiata*, we searched a graph of the laser-scanned foliage that allowed both walking on the surface and jumping across gaps. Mesh vertices were merged onto a 2 mm grid to form the graph's nodes. Two nodes were connected by a walking edge if the mesh connected them, with a cost equal to the distance between them. Connections came from the mesh rather than from proximity, so surfaces that came close without touching, such as two overlapping leaves, stayed separate and could only be crossed by a jump. Jumps were allowed only between separate surfaces. From each surface point, we traced rays within a cone of half-angle 35° around the surface normal; a ray that struck another surface within 3.5 cm defined a possible jump. The 35° limit is an assumption, chosen to exclude jumps nearly parallel to a surface, rather than a measured value. A* search[70] (implemented in Python's NetworkX package) was used to solve for the shortest path from *P. labiata*'s reroute position to the prey spider's position.

For analyzing *P. labiata*'s habitats in general, four other regions of foliage where *P. labiata* were found were scanned for lacunarity and geometric analysis. Lacunarity is a measure of how 'gappy' a space is, and has been adopted by land ecologists. High lacunarity corresponds to large, clumped gaps; low lacunarity corresponds to small, evenly spread gaps[71, 72]. We computed the 3D lacunarity[73] with a custom Python script.

We also computed the vision-based network degree complexity as our measure of spatial complexity[16, 50] within the habitat environments. For the hunt itself, we used the 12 cm radius volume from the prey's position: for the other four habitat laser scans, we used a 12 cm radius as well, chosen to avoid edge effects. Foliage within these volumes was voxelized using Trimesh (4.11.5). Regions were voxelized by rounding the (subdivided) scan surface onto a 6 mm grid, marking a voxel occupied if the surface passes within half a voxel of its center. For these volumes we sampled the possible voxels that *P .labiata* could perch on (the open-air voxels that are adjacent to surface voxels) which places each viewpoint one voxel clear of the foliage. This set numbered 4.7–7.6 $\times 10^4$ voxels per region. 40,000 of these voxels were uniformly randomly sampled and used as nodes (corresponding to 52–84% of the perch point voxels) of the habitats. An edge then joins two nodes unless they are obstructed by another object. The network degree complexity is given by $C$

$$C = \frac{H}{\log D} \tag{2}$$

The denominator $\log D$ is the entropy of a uniform distribution over the D attainable degrees, and

$$H = -\sum_{d=1}^{D} f(d) \log f(d) \tag{3}$$

$$\text{where } f(d) = \frac{n_d}{D}, \qquad d = 1, \ldots, D \tag{4}$$

with $n_d$ as the number of nodes of degree $d$, and $D$ the total number of nodes. $C$ ranges from 0 to 1. $C = 0$ when every node has the same degree, and $C = 1$ when every node has a different degree. Our wild hunt location reported a vision-based network complexity of 0.87, implying a higher degree of complexity to that of a sample of savanna (described in caption of Supplementary Fig. 2), at 0.80[50]. We report the vision-based network complexities of the 5 habitat laser scans in Supplementary Table 1.

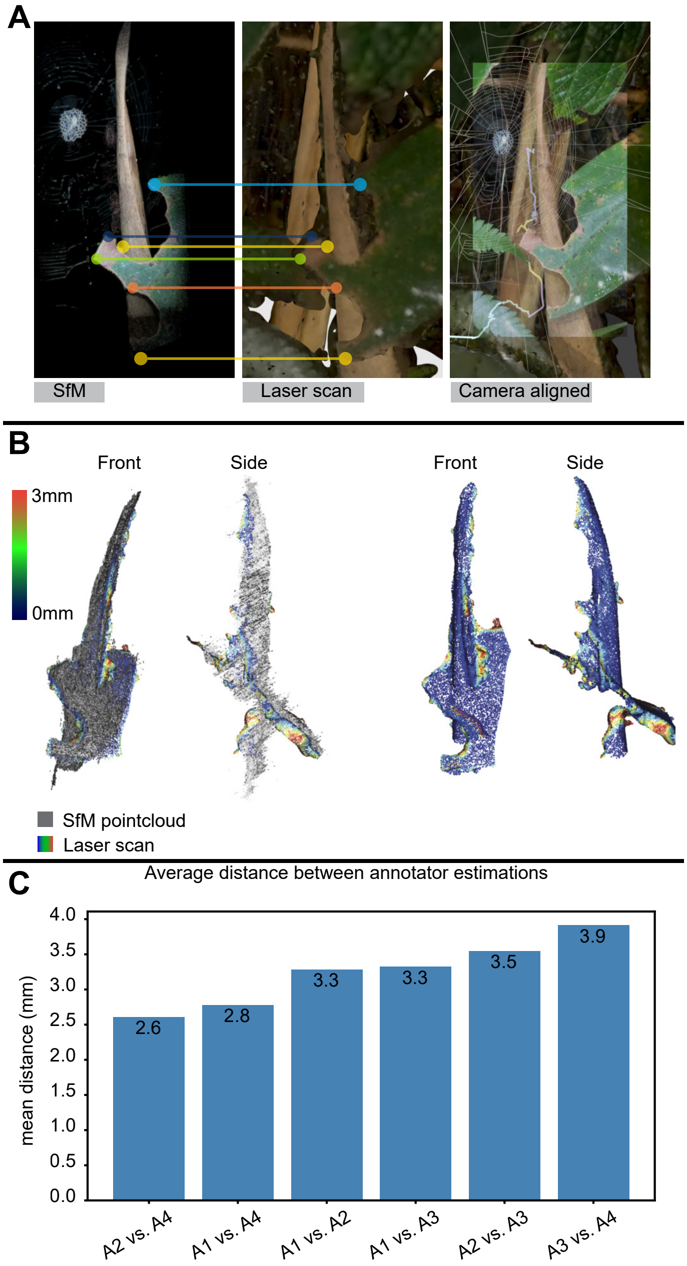


Supplementary Figure 1: **A.** Alignment of the SfM point cloud (left) to the laser scan (middle) using the corresponding features shown as colored dots; on the right, an iPhone 12 Pro frame overlaid on the laser scan shows the result. The alignment gives the reconstruction metric scale. **B.** Alignment error: distance from each laser-scan point to the closest point in the aligned SfM point cloud (heat map; SfM in grey), front and side views. Median 0.41 mm (interquartile range 0.12–1.22 mm; mean ± SD 0.88 ± 1.13 mm); the largest errors are where SfM did not reconstruct the surface. **C.** Mean position error between manual annotators across all tracked points and frames: 2.6–3.9 mm, about half of *P. labiata*'s body width.

Supplementary Table 1: Network degree complexity $C$ of sight lines of the five habitat regions.

| Region | $C$ |
|---|---|
| Location 1 | 0.894 |
| Location 2 | 0.875 |
| Hunt location | 0.873 |
| Location 5 | 0.851 |
| Location 4 | 0.837 |

Supplementary Table 2: Software used for reconstruction and alignment, with versions and the settings reported in Methods. Software for tracking and analysis: Supplementary Table 3.

| **Software, version** | **Usage** | **Settings** |
|---|---|---|
| Artec Studio Professional 18.1.4.9 | Laser scans (Artec Leo) of the hunt site and four other habitats: registration, cleanup and mesh fusion | *Scanning:* geometry and texture, HD mode, high resolution, 40 frames $s^{-1}$; 9 million polygons (8 million, location 2). *Global registration:* keyframe ratio 0.4, then 0.6. *Fusion resolution:* 0.5–0.7 mm. *Hunt-site model:* 4,493,317 vertices, 2 million polygons |
| Artec Studio 19 (beta) | Photogrammetry of a female *P. labiata* (not the hunting individual) | *Photographs:* POCO X6 Pro rear camera, flash off, inside a PULUZ light box |
| COLMAP 3.12.3[60], GLOMAP 1.2.0[61] | SfM of both hunt videos (iPhone 12 Pro, 1080 × 1920, 59.97 fps; iPhone X, 1920 × 1080, 30 fps) | *Features:* SIFT, 4 octaves, edge threshold 10.00; `SIMPLE_RADIAL`, one camera per video. *Matching:* exhaustive. *Sparse:* COLMAP incremental, 913 frames (iPhone 12 Pro, downsampled to 30 fps); GLOMAP global, 2,375 frames (iPhone X). *Dense:* COLMAP, default settings. *Registration:* `image_registrator` at 5 Hz; 5,476 (12 Pro), 12,922 (X) frames; the two reconstructions then merged into one point cloud |
| GLOMAP 1.2.0[61], COLMAP 3.12.3[60] | SfM of the final capture (iPhone 12 Pro), for the full-pose reconstruction | *Input:* 2,678 frames (every 9th); features as above, one camera. *Matching:* 12 neighbours either side, plus every 10th frame against each other (67,569 pairs). *Sparse:* GLOMAP global. *Refinement:* COLMAP bundle adjustment, focal length and $k_1$ refined. *Result:* 2,671 frames registered, 77,803 points, 1.21 px reprojection error |
| Open3D 0.19.0[65] | Scaling and aligning the SfM point cloud to the laser scan (hunt videos) | *Initial fit:* Procrustes[63] on 6 corresponding features, then manual adjustment. *Refinement:* ICP[64]. *Error:* median distance from each laser-scan point to the nearest aligned SfM point 0.41 mm (interquartile range 0.12–1.22 mm; mean 0.88 ± 1.13 mm) |
| Python (OpenCV, SciPy; custom script) | Additional refinement of both cameras (final-capture clip, iPhone X) against the modelled stabilimentum | *1. Silk:* white top-hat (15 px) of the brightness, kept where saturation $\leq 0.35$ and within 220 px of the projected stabilimentum; distance from every pixel to the nearest silk. *2. Model:* points every ≈2 px along the modelled stabilimentum's edges, projected through each frame's camera. *3. Fit:* image-shift search (±160 px, 4 px steps), then least squares on a small camera rotation, distances capped at 12 px; camera distance held. *4. Quality:* frames kept if ≥35% (clip) or ≥40% (iPhone X) of points lie within 3 px of silk; clip sampled every 9th frame, before the strike only. *5. Smoothing:* median over 7 samples, Gaussian ($\sigma = 2$ samples), interpolated to every frame |
| Python 3.13; Blender 5.1 (custom scripts) | Web plane, strands and anchor points | *RANSAC*[62]: on a manually picked web region of the point cloud; inlier threshold 0.2, 1000 iterations; plane, then strands. *Missing strands:* drawn in the view plane of iPhone 12 Pro frames and projected onto the web plane. *Anchors:* first foliage intersection along each radial's direction, by ray casting against a bounding volume hierarchy (BVH) of the laser-scan mesh (Blender `mathutils.bvhtree`) |
| OpenCV 4.12 | Enhancing silk in video frames for tracing | *Gaussian smoothing:* $\sigma = 1$ px. *White top-hat:* elliptical, 15 px. *Contrast stretch:* 85th–99.8th percentile. *Gamma:* 0.7. *Overlay:* pale cyan, on the frame 45% darker and 60% desaturated |
| SuperPoint, LightGlue[76, 77]; pycolmap 3.10 (custom script) | Registering the Nikon COOLPIX P950 photographs into the aligned iPhone 12 Pro reconstruction | *Matching:* each photograph matched to every 40th video frame, then to the 8 best and their neighbours (4096 keypoints; RANSAC-filtered). *Pose:* absolute pose from the matched 3D points, refining focal length and $k_1$; ≥40 inliers. *Result:* 17 of 18 photographs registered |
| Python (OpenCV, custom script) | Silk in the registered photographs, combined on the web plane | *Threads:* high-pass, dew drops inpainted, two-sided line averaging at 36 orientations, normalized by local noise; contrast-stretched between the 90th and 99.8th percentiles, gamma 0.6. *Combination:* each photograph projected onto the web plane; share of the photographs seeing a point that find silk there |

Supplementary Table 3: Software used for tracking and analysis, with versions and the settings reported in Methods. Software for reconstruction and alignment: Supplementary Table 2.

| **Software, version** | **Usage** | **Settings** |
|---|---|---|
| DeepLabCut 2.3[66, 67] | Tracking body (whole hunt) and legs (final capture) | *Keypoints:* cephalothorax (between the anterior median eyes) and abdomen centre; in the final capture also coxa, tibia and tarsus of each leg (26). *Labelled frames:* 500 per model. *Likelihood cutoff:* $\geq 0.99$ |
| Python 3.13 (custom script) | Solving spider position and pose from tracked rays, with manual correction | *One view:* least-squares convergence of rays from adjacent frames. *Two views:* ray intersection (stalk sequence). *Final capture:* body by ray convergence; IK for legs on the leaf, FK along coxa–tibia–tarsus for legs in the air. *Manual:* 286 frames corrected and oriented by 4 annotators (157 by annotation alone; inter-annotator error 2.6–3.9 mm); 16 final-capture frames oriented by 4 annotators |
| SciPy 1.17[68] | Speed smoothing (`signal.savgol_filter`) | *Filter:* Savitzky–Golay, non-causal; window 31 s (31 samples on a 1 s grid); polynomial order 3. *Not captured:* hesitant jump and final attack, estimated instead from displacement over 3 consecutive frames (450 and 600 mm $s^{-1}$) |
| Blender 5.1 | Rendering; measurement; leg gait (custom script) | *Renderer:* Cycles path tracer. *Measurement:* prey ≈6 mm (front to rear leg tips), discoid ≈9 mm diameter. *Gait:* body follows the 3D trajectory; IK on 8 foot targets |
| Blender 5.1 Python (custom script) | Visual fields and line of sight: prey to *P. labiata* and its routes; *P. labiata* to the prey | *Prey:* eye fields from Land and Barth's maps[25], deprojected with the Hammer projection; ranges 9.9, 7.9, 31.8, 28.6 cm (AM, AL, PM, PL) for a 10 mm target; 40° below the web plane. *Occlusion:* ray casting against a bounding volume hierarchy (BVH) of the laser-scan mesh and the prey's discoid (Blender `mathutils.bvhtree`); a point is hidden if the ray to the prey's eyes hits either first. *P. labiata:* 10.5 cpd; retinal tubes ±50° horizontal, 35° vertical, plus ≈30° body roll and pitch. *Routes:* 10 mm tube; seen if visible in any frame every 500 frames (≈32 s), frames 0–8254 |
| Python (custom script) | 3D lacunarity[73] of five habitats | *Boxes:* normalized by *P. labiata*'s body length. *Lacunarity value:* area under the curve[75] |
| NetworkX (version not stated) | A* shortest path[70] from the reroute position to the prey | *Nodes:* mesh vertices merged onto a 2 mm grid. *Walk edges:* where the mesh connects nodes, cost = length. *Jumps:* 32 rays per surface point, spread evenly within a cone of half-angle 35° around the surface normal (an assumed limit) and ray marched; the first surface each reaches within 3.5 cm is a possible landing, cost = straight-line distance |
| Trimesh 4.11.5 | Voxelization for vision-based network degree complexity[16, 50] | *Voxels:* 6 mm; occupied if the subdivided scan surface passes within half a voxel of the centre. *Region:* 12 cm radius around the prey (other habitats: 12 cm). *Nodes:* 40,000 of the 4.7–7.6 × $10^4$ perch voxels (52–84%), joined unless the line of sight is blocked. *Result:* $C = 0.837$–0.894; hunt location 0.873 |

Supplementary Table 4: Vision models

| | Prey's view (*A. versicolor*) | *P. labiata*'s view |
|---|---|---|
| Acuity basis | *C. salei* eyes[25], used as an upper bound because wandering spiders see better than orb weavers; 0.139 cpd to 0.56 cpd | *P. fimbriata* measured at 12.5 cpd[20], scaled by lens size[46] to ≈10.5 cpd |
| Target model | *P. labiata* taken as 10 mm (upper bound of 6 mm to 10 mm) | Prey measured at ≈6 mm; routes modeled as a 10 mm tube over the foliage surface |
| Range | 7.9 cm to 31.8 cm (by eye pair) | 3.6 m; maximum predator–prey distance during the hunt was 11 cm |
| Field of view | Eye maps of *C. salei*[25] and permitting 40 degrees FOV beneath the web plane. Line of sight tested against the discoid derived from SfM, and the laser scan foliage. | Retinal tubes move 50° horizontally and 35° vertically[47], plus ≈30° body roll and pitch |
| Eye position | Placed on the head of *A. versicolor* derived from the SfM reconstruction. Does not move throughout hunt. | Cephalothorax tracked with DLC across the videos. 7,642 frames were used to solve *P. labiata*'s position across the stalk sequence. 540 frames were used to solve for positions in the first half of the hunt. 286 frames were taken across the hunt and manually oriented as well as corrected for if necessary. |
| What limits visibility? | Range, field of view, and occlusion | Field of view and occlusion. |
| Direction of bias of visual model | Overestimates what the prey sees using a better model. Eye angles not known for *A. versicolor*; they have a fixed FOV[78, 79] | Overestimates what *P. labiata* was focused on. Unknown where *P. labiata* is focusing its fovea, thus visual model takes the entire possible visual field. |

Supplementary Table 5: Route comparison

| Question | Window and position | Measurement | Against shortest path | Against taken path |
|---|---|---|---|---|
| Was *P. labiata* visible to the prey on this route (post-aggressive mimicry)? | Prey, whole route | % of route length inside the prey's visual volume | $\approx 97\%$ | $\approx 41\%$ |
| Could *P. labiata* see this route before choosing? | Start to reroute | % of route length inside *P. labiata*'s visual field (upper bound: 50° horizontal, 35° vertical, allowing ±30° body pitch and roll) | 77%–100% | $\leq 80\%$ |
| Could *P. labiata* see the prey? | Position and orientation over the hunt | Time and length of the path with the prey not visible | 58% of total path length | 40% of total path length; 24% of hunt duration |
| How many web guylines were connected to foliage along the path? | Total path | Total number of silk connections over surfaces that the path crosses | 10–12[a] | 1–3[a] |

[a]Upper values allow the web to shift by the maximum possible depth error, in which case more guylines connect to the leaf that the shortest path crosses, and the web connects to the top of the right brown leaf that *P. labiata* stalked over.

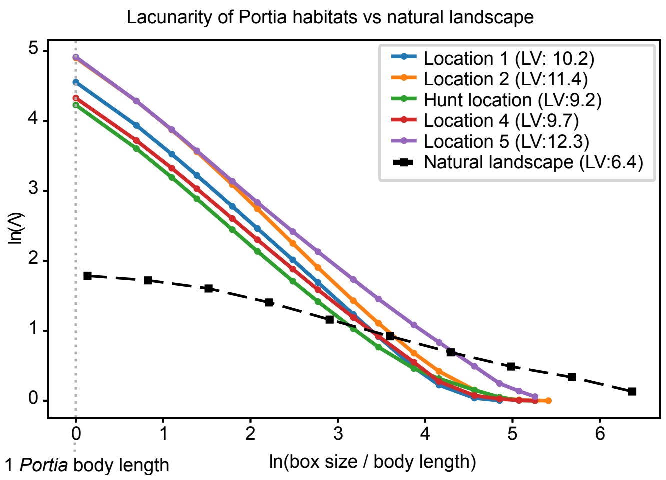


Supplementary Figure 2: Lacunarity curves of five *P. labiata* habitats (including the hunt domain, details: Methods). Boxes are normalized by *P. labiata*'s body length; for the natural environment (previously published[50]: Supplementary Fig. 1) boxes are normalized by average human body length (approximately 1.75 m). The landscape is a binarized Google Earth image of savanna in the Okavango Delta, Botswana, 2730 × 1689 pixels covering 1841 m × 1139 m, so each pixel covers 0.674 m × 0.673 m (0.45 $m^2$). The curves decline linearly, which is typical of self-similar patterns[74]. Lacunarity value (LV): area under the curve[75].
